# *In vivo* effects of inducible transgene expression of the age-associated glycogenes *St6gal1* and *Man1a* on epidermal homeostasis

**DOI:** 10.64898/2026.11.03.763605

**Authors:** Trisha Biswas Shanta, Nguyen Thi Kim Nguyen, Erna Raja, Mizuho Ishikawa, Ahmed G. K. Habib, Lalhaba Oinam, Hiromi Yanagisawa, Hiroaki Tateno, Aiko Sada

## Abstract

Glycosylation of cell-surface proteins plays several essential roles in cell–matrix interactions and receptor signaling, thereby contributing to tissue homeostasis and aging. In aged mouse skin, epidermal stem cells exhibit increased levels of α2,6-sialylated complex-type N-glycans alongside upregulation of glycogenes such as *St6gal1* and *Man1a*; however, the functional importance of these glycosylation changes *in vivo* remains largely unknown. Here, we generated doxycycline-inducible transgenic mice expressing the age-related glycogenes *St6gal1* and *Man1a,* either individually or in combination, using ubiquitous and epidermis-specific drivers. Lectin-based analyses with SNA and rHeltuba demonstrated corresponding alterations in glycosylation patterns in these mice. Transgenic *Man1a* induction reduced epidermal proliferation and epidermal thickness. In contrast, *St6gal1* induction caused splenomegaly and elevated expression of inflammatory signature genes in the skin but did not measurably alter epidermal proliferation or morphology. Double transgenic mice exhibited a hybrid phenotype combining inflammatory and proliferative impairments, although comparison of ubiquitous and epidermis-specific induction indicated that the observed skin phenotypes depended on the epidermal expression of the transgenes. An SNA lectin-based pull-down assay followed by mass spectrometry identified candidate α2,6-sialylated proteins, including integrin family members, junctional proteins, and surface receptors, following induction of *St6gal1* and *Man1a*. Together, these findings suggest that increased expression of the age-associated glycogenes *St6gal1* and *Man1a* perturbs distinct aspects of epidermal homeostasis *in vivo* and identify candidate glycoproteins that may mediate these effects.

**Graphical Abstract:** 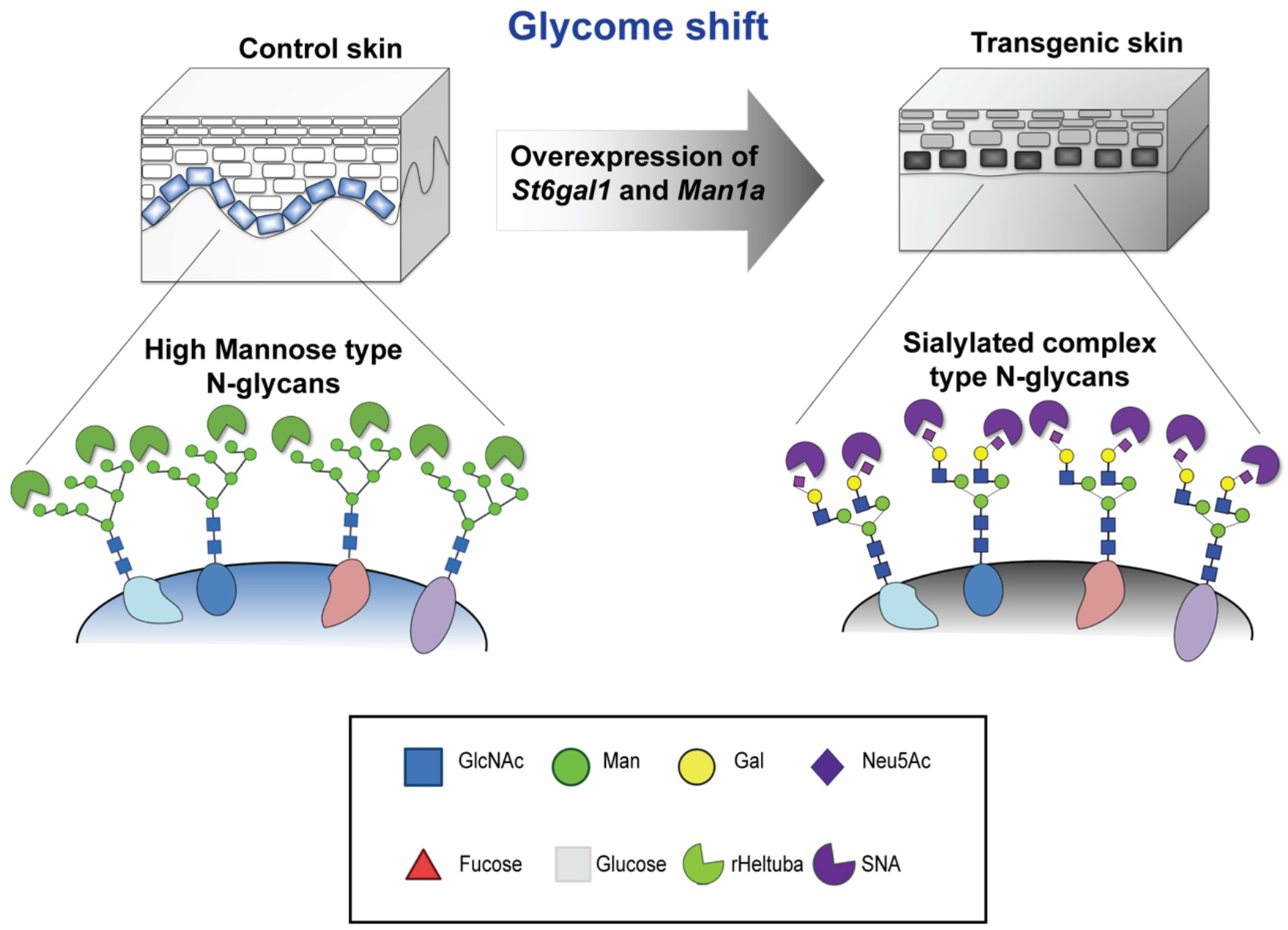

## Introduction

Aging is accompanied by a progressive decline in tissue function, driven by cellular and molecular processes collectively described as the hallmarks of aging (López-Otín et al. 2023). Although transcriptional, epigenetic, and metabolic alterations associated with aging have been widely studied, post-translational modifications—particularly glycosylation—remain underexplored. Glycans are complex sugar chains attached to proteins and lipids that regulate cell–cell communication, stem cell function, receptor signaling, and extracellular matrix (ECM) interactions (Varki 2017; Reily et al. 2019; Broussard & Boyce 2019; Nishihara 2018). However, how age-associated changes in glycosylation influence tissue homeostasis *in vivo* remains unclear.

The skin epidermis is continuously renewed by epidermal stem cells that maintain tissue architecture and barrier function. Aging impairs epidermal homeostasis and is associated with epidermal thinning, reduced proliferative capacity, impaired wound healing, and increased inflammation (Gilchrest 1983; Changarathil et al. 2019; Keyes et al. 2016; Giangreco et al. 2008). These age-dependent changes arise not only from cell-intrinsic factors but also from altered interactions between epidermal stem cells and their niche microenvironment. In particular, cell-surface receptors and adhesion molecules, including integrins, are known to be highly glycosylated and play important roles in epidermal stem cell regulation and age-associated tissue dysfunction (Rao et al. 2022; Ankenbauer et al. 2023; Liu et al. 2024).

Glycosylation has emerged as an important regulator of cellular function and disease in the skin (Shen et al. 2025). CRISPR/Cas9-mediated knockout of glycosyltransferases in human skin culture models has been shown to impair epidermal barrier formation and differentiation (Dabelsteen et al. 2020). Glycomic analysis has revealed that the murine epidermis is enriched in high-mannose glycans, but shows less abundant sialylation and fucosylation compared with the dermis (Uematsu et al. 2009). Using a lectin microarray (Tateno et al. 2011; Wang et al. 2015), we previously identified an aging-associated glycome shift in epidermal stem cells, characterized by increased α2,6- and α2,3-sialylation (Oinam et al. 2020). Similar glycosylation patterns and age-associated changes have also been observed in human skin (Möginger et al. 2018; Abdayem et al. 2016; Donovan et al. 2023). Among the glycogenes associated with these changes, *St6gal1* and *Man1a* were upregulated in epidermal stem cells during aging, and functional assays of glycogenes in cultured keratinocytes further suggested that these enzymes influence the proliferation ability of these cells (Oinam et al. 2020). *Man1a* functions as a Golgi-resident α-mannosidase that trims high-mannose residues, while *St6gal1* catalyzes the addition of α2,6-linked sialic acid to N-glycans (Figure 1a) (Kornfeld & Kornfeld 1985; Togayachi et al. 2026; Schjoldager et al. 2020; Moremen et al. 2012). Although both enzymes catalyze key N-glycan maturation reactions, their functional significance in maintaining epidermal homeostasis has not been defined.

**Figure 1.**
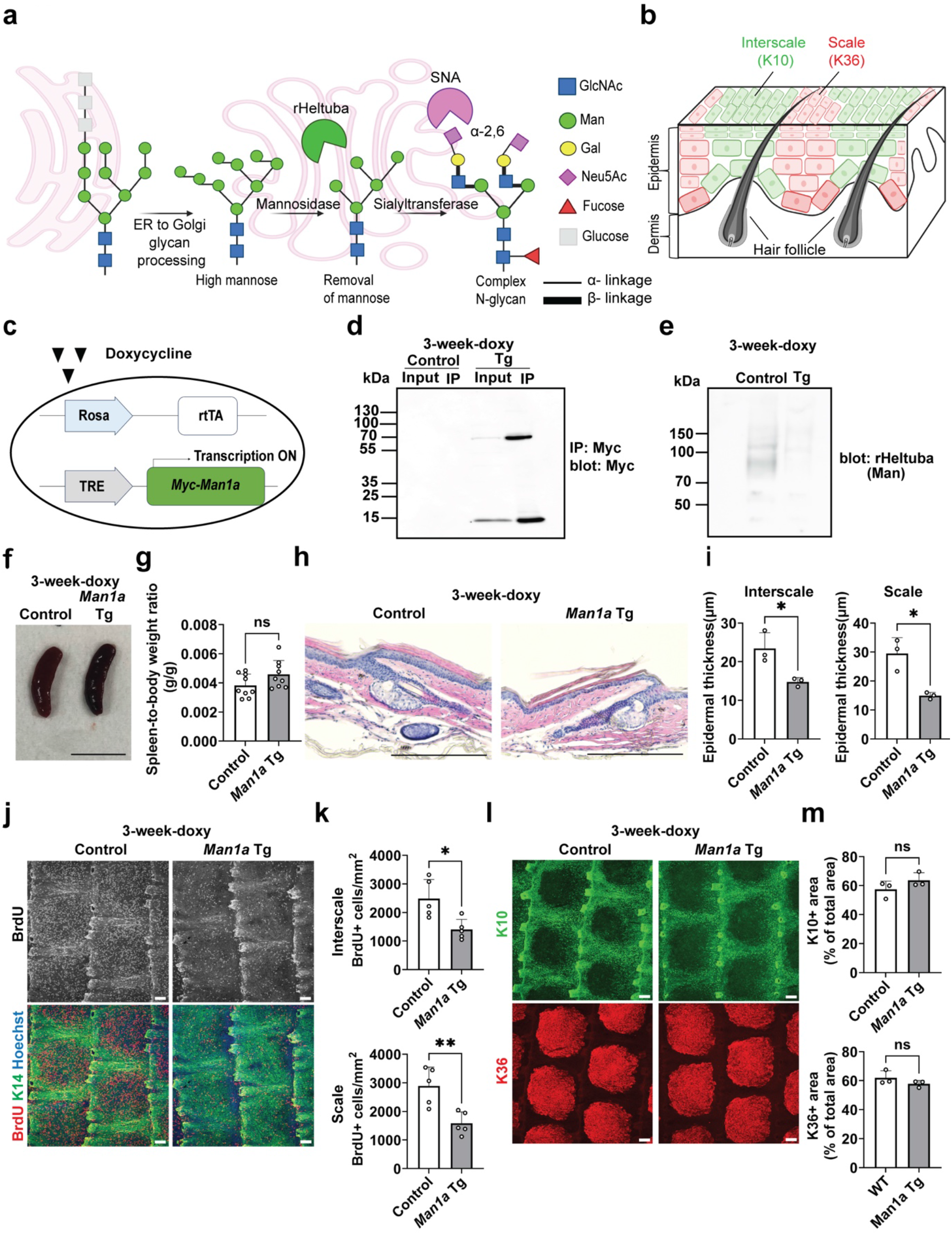
*Man1a* induction leads to decreased epidermal thickness and proliferation. (**a**) Schematic representation of the N-glycosylation process. (**b**) Schematic representation of the mouse tail skin. (**c**) Doxycycline-inducible Tet-On system used to express Myc-tagged *Man1a*. (**d**) Detection of Myc-tagged *Man1a* by anti-Myc immunoprecipitation. (**e**) rHeltuba (a mannose binder) blotting of membrane protein fractions from FACS-isolated epidermal basal cells. (**f, g**) Representative spleen picture and spleen/body weight ratio from control and *Man1a* Tg mice. Scale bar, 10 mm. (**h, i**) Hematoxylin and eosin (H&E) staining of tail skin (h) and quantification of epidermal thickness (i) after 3 weeks of doxycycline treatment in control and *Man1a* Tg mice. Scale bars, 300 µm. (**j, k**) Whole-mount staining of tail epidermis for BrdU (proliferation, red), K14 (epidermal basal cells, green), and Hoechst (blue) (j) and quantification of BrdU-positive epidermal cells (k). Scale bars, 100 µm. (**l, m**) Whole-mount staining for K10 (interscale, green), K36 (scale, red), and Hoechst (blue) (l) and quantification of K10- and K36-positive areas (m). Scale bars, 100 µm. Data are presented as the mean ± SD. Each dot represents one mouse, and measurements from multiple regions of the same mouse are averaged. Comparisons between control and Tg groups were performed using two-tailed unpaired Student’s t-tests. \**P* < 0.05; \*\**P* < 0.01; ns, not significant.

α2,6-Sialylation mediated by *St6gal1* has been implicated in diverse physiological and pathological processes (Zhu et al. 2024). Studies have shown that *St6gal1* modulates inflammatory responses, including the regulation of cytokine production and eosinophilic inflammation (Krick et al. 2021; Yasukawa et al. 2005; Nasirikenari et al. 2010; Wang et al. 2021). In addition, age-related remodeling of sialylated glycans in serum and solid tissue has been reported (Hanisch et al. 2013; Catera et al. 2016; Zhang et al. 2025). Dermal photoaging is also induced by the upregulation of *St6gal1* (Zhong et al. 2026). Mechanistically, *St6gal1* modifies surface receptors, such as EGFR, TNFR1, and integrins, thereby influencing receptor stability and downstream growth factor and immune signaling (Kaburagi et al. 2017; Liu et al. 2024b; Gu & Taniguchi 2004; Holdbrooks et al. 2020; Holdbrooks et al. 2018). In contrast, the functional roles of *Man1a* have been less widely characterized. Clinical and experimental studies suggest that *Man1a* contributes to tumorigenesis in a context-dependent manner: upregulation of *Man1a* promotes proliferation and activates the unfolded protein response in hepatocellular carcinoma (Tu et al. 2017), whereas reduced *Man1a* expression correlates with aberrant N-glycosylation and poor prognosis in breast cancer (Legler et al. 2018).

Together, these previous studies raise the possibility that age-associated upregulation of *St6gal1* and *Man1a* may alter N-glycan processing and thereby affect epidermal function during aging. However, the consequences of such glycan remodeling for epidermal homeostasis *in vivo* remain unclear. In this study, we generated doxycycline-inducible transgenic mouse models expressing *St6gal1* and *Man1a in vivo*. Increased expression of these glycogenes perturbed distinct aspects of epidermal homeostasis, including epidermal proliferation, tissue morphology, and inflammatory gene expression. Lectin-based glycoproteomic analysis further identified candidate α2,6-sialylated membrane proteins, including integrins and other cell-surface proteins. Together, our findings support a potential link between age-associated N-glycan remodeling and epidermal tissue dysfunction.

## Results

### *Man1a* induction reduces epidermal proliferation and thickness

N-glycosylation proceeds sequentially, with *Man1a* trimming high-mannose precursors and *St6gal1* then catalyzing terminal α2,6-sialylation (Schjoldager et al. 2020; Moremen et al. 2012) (Figure 1a). The mouse tail epidermis is spatially organized into K10^+^ interscale and K36^+^ scale compartments (Gomez et al. 2013), providing a system in which epidermal proliferation, thickness, lineage organization, and their age-related changes can be evaluated (Figure 1b). Previous work has demonstrated that aging skin exhibits altered glycosylation patterns, including increased α2,6-sialylation and upregulation of the glycogenes *St6gal1* and *Man1a* (Oinam et al. 2020). To investigate their functions *in vivo*, we generated doxycycline-inducible Tet-On transgenic (Tg) mice, in which each gene was expressed under the control of a pTRE promoter (Figure 1c). These mice were then crossed with either Rosa-rtTA mice for ubiquitous induction or K14-rtTA mice for epidermis-specific induction. Upon doxycycline administration, rtTA activates the pTRE promoter, enabling temporal and tissue-dependent expression of *St6gal1* and *Man1a*.

In order to validate transgene induction, young adult mice were fed doxycycline-containing chow for 3 weeks. Immunoprecipitation followed by western blotting was used to detect Myc-tagged *Man1a* in epidermal lysates from *Man1a* Tg mice (Figure 1d). To assess the enzymatic activity of the transgene, epidermal basal cells were isolated by FACS, and membrane proteins were then subjected to lectin blotting using rHeltuba, a lectin probe that preferentially binds to mannose. *Man1a* Tg mice exhibited a decrease in rHeltuba binding (Figure 1e), indicating functional enzymatic activity.

To address how *Man1a* induction affects epidermal homeostasis, we analyzed epidermal structure, epidermal stem cell proliferation, and differentiation lineages following doxycycline induction in *Man1a* Tg mice. After 3 weeks of doxycycline treatment, *Man1a* Tg mice did not develop splenomegaly, suggesting an absence of systemic inflammation (Figure 1f, g). Histological analysis of *Man1a* Tg skin revealed a significant reduction in epidermal thickness (Figure 1h, i). BrdU incorporation assays showed a significant decrease in epidermal proliferation (Figure 1j, k), consistent with the growth defects caused by *Man1a* induction *in vitro* (Oinam et al. 2020). Despite these changes, K10/K36 lineage marker analysis showed predominant preservation of the interscale/scale organization (Figure 1l, m), which has been reported to be disrupted in aging skin (Changarathil et al. 2019; Raja et al. 2022; Fan et al. 2026). These results suggest that elevated *Man1a* expression primarily impairs epidermal proliferation and morphology.

### *St6gal1* induction produces inflammatory features but preserves epidermal tissue architecture

Next, we examined the phenotypes of *St6gal1* Tg mice (Figure 2a). After 3 weeks of doxycycline treatment, Flag-tagged *St6gal1* was detected in epidermal lysates by Flag-based immunoprecipitation followed by western blotting (Figure 2b). Lectin blotting with SNA, a lectin probe that preferentially binds to terminal α2,6-sialic acids (Shibuya et al. 1987), further revealed increased SNA binding in epidermal basal cells of *St6gal1* Tg mice (Figure 2c). In contrast to *Man1a* Tg mice, *St6gal1* Tg mice developed splenomegaly after 6 weeks of induction (Figure 2d, e), and showed no appreciable changes in epidermal thickness or overall skin morphology (Figure 2f, g). BrdU incorporation assays revealed comparable levels of epidermal proliferation between Tg and wild-type controls (Figure 2h, i). Whole-mount staining with epidermal differentiation markers (K10/K36) further demonstrated preserved interscale/scale organization (Figure 2j, k). These results indicate that *St6gal1* induction does not measurably alter epidermal proliferation, structure, or interscale/scale organization, in contrast to the phenotypes observed in *Man1a* Tg mice.

**Figure 2.**
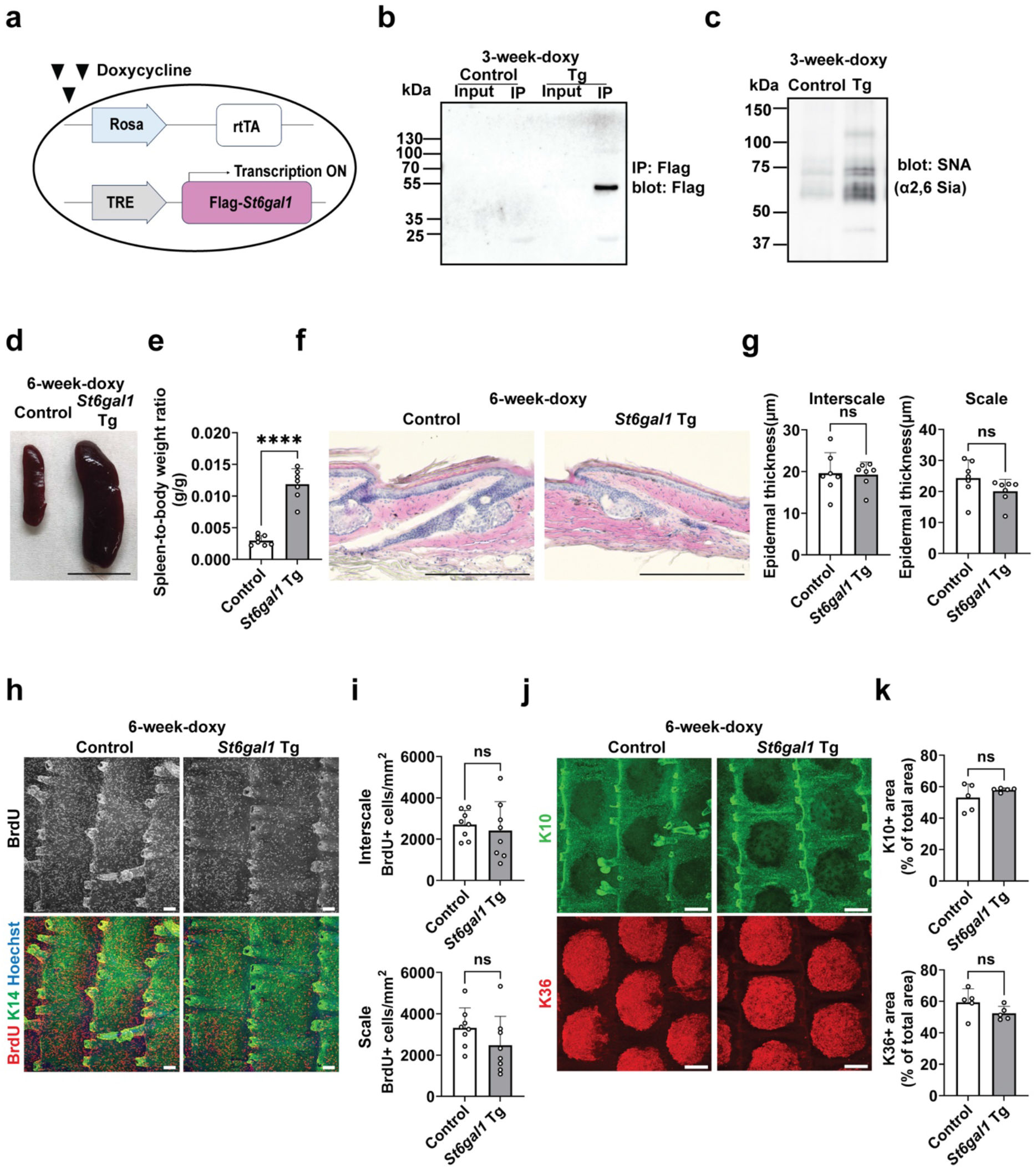
Validation and phenotypic analysis of doxycycline-inducible *St6gal1* Tg mice. (**a**) Doxycycline-inducible Tet-On system used to express Flag-tagged *St6gal1*. (**b**) Detection of Flag-tagged *St6gal1* by anti-Flag immunoprecipitation. (**c**) SNA (an α2,6-sialic acid binder) lectin blotting of membrane protein fractions from FACS-isolated epidermal basal cells. (**d, e**) Representative spleen picture and spleen/body weight ratio from control and *St6gal1* Tg mice. Scale bar, 10 mm. (**f, g**) H&E staining of tail skin (f) and quantification of epidermal thickness (g) from control and *St6gal1* Tg mice after 6 weeks of doxycycline treatment. Scale bars, 300 µm. (**h, i**) Whole-mount staining of tail epidermis for BrdU (proliferation, red), K14 (epidermal basal cells, green), and Hoechst (blue). Scale bars, 100 µm. Quantification of BrdU-positive epidermal cells (i). (**j, k**) Whole-mount staining for K10 (interscale, green), K36 (scale, red), and Hoechst (blue). Scale bars, 200 µm. Quantification of K10- and K36-positive areas (k). Data are presented as the mean ± SD. Each dot represents one mouse, and measurements from multiple regions of the same mouse are averaged. Comparisons between control and Tg groups were performed using two-tailed unpaired Student’s t-tests. \*\*\*\**P* < 0.0001; ns, not significant.

To examine the molecular changes induced by *St6gal1* expression, we performed bulk RNA-seq on epidermal basal cell populations (integrin α6^high^/CD34^−^/Sca-1^+^) isolated from *St6gal1* Tg/Rosa-rtTA mice after 6 weeks of doxycycline treatment (Figure 3a). Principal component analysis (PCA) and hierarchical clustering revealed a separation of *St6gal1* Tg and control groups, indicating their distinct molecular properties (Figure 3b, Supplemental Table S1). Gene Ontology (GO) analysis showed enrichment of downregulated genes in pathways related to epidermal homeostasis, including intermediate filament organization, the tyrosine kinase signaling pathway, and ECM organization (Figure 3c). In contrast, upregulated genes were enriched in inflammatory and pathological remodeling pathways, including cytokine, interferon, and TNF signaling (Figure 3d). Consistently, gene set enrichment analysis (GSEA) and heatmaps showed trends of negative enrichment in gene sets associated with epidermal morphogenesis and cell–cell adhesion, and trends of positive enrichment for type II interferon response and Toll-like receptor signaling gene sets (Figure 3e, f). Thus, systemic *St6gal1* induction was associated with an inflammatory transcriptional signature and reduced expression of epidermal homeostasis-related genes in epidermal basal cells, despite the absence of overt histological abnormalities.

**Figure 3.**
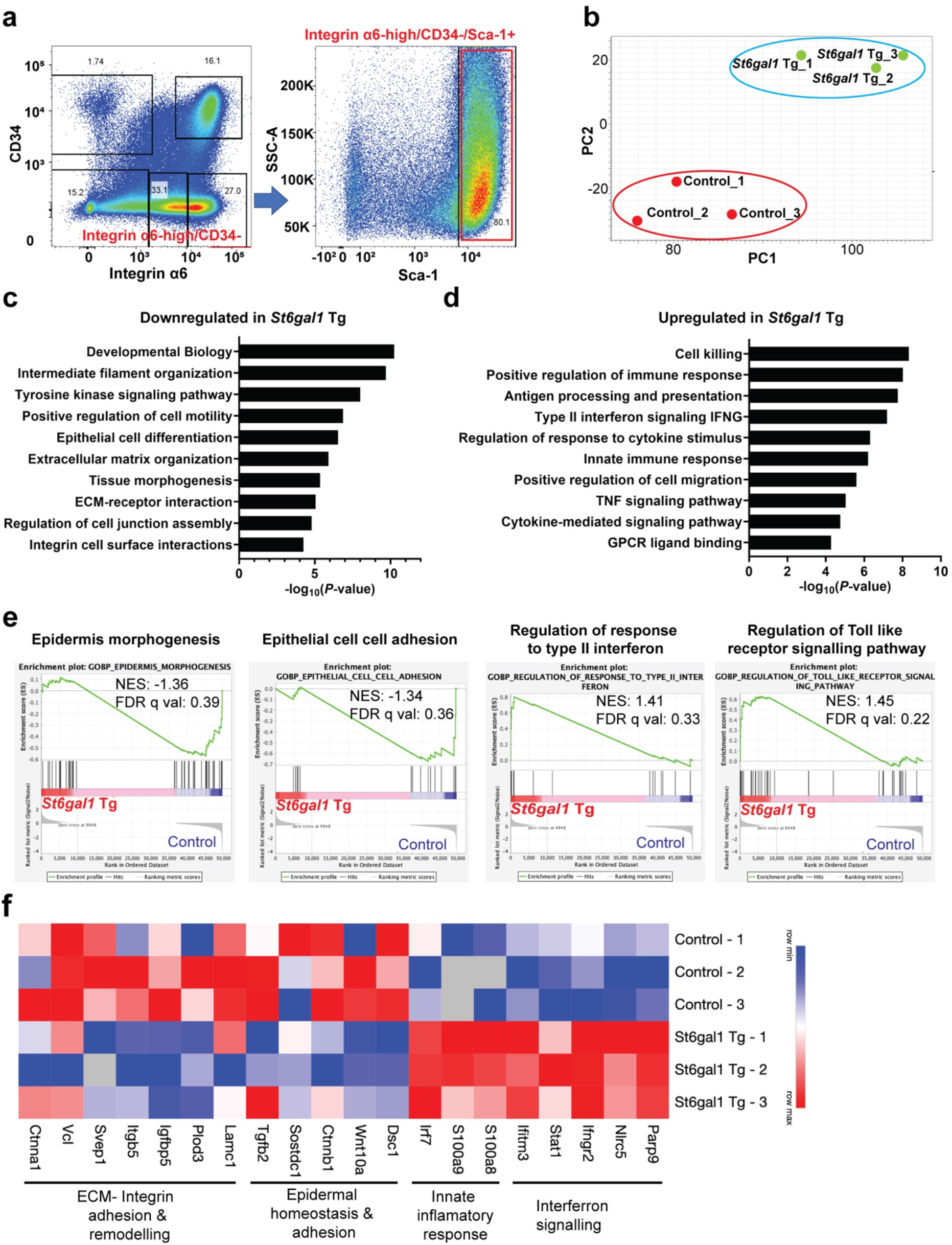
*St6gal1* induction is associated with inflammation-related transcriptional changes in epidermal basal cells. **(a)** Representative FACS plot showing the isolation of viable integrin α6^high^/CD34^−^/Sca-1^+^ epidermal basal cells from control and *St6gal1* Tg mice after 6 weeks of doxycycline treatment. (b) Principal component analysis of RNA-sequencing data from FACS-isolated epidermal basal cells (n = 3 mice per group). (**c, d**) Gene Ontology analysis of differentially expressed genes showing at least a twofold downregulation (c) or upregulation (d) in *St6gal1* Tg relative to control mice. (**e**) Gene set enrichment analysis (GSEA) showing positively enriched inflammation-related gene sets and negatively enriched gene sets associated with epidermal function in *St6gal1* Tg mice. (**f**) Heatmap showing the relative expression of selected leading-edge genes from GSEA-enriched gene sets associated with inflammatory and interferon responses, integrin–ECM interactions, and epidermal homeostasis. Each row represents one mouse, and each column represents one gene.

### Combined *St6gal1* and *Man1a* expression produces systemic and epidermal phenotypes in a tissue-dependent manner

To determine whether *St6gal1* and *Man1a* act synergistically or independently, we generated double transgenic mice expressing both glycogenes, by using either a ubiquitous Rosa-rtTA driver (Figure 4a) or an epidermis-specific K14-rtTA driver (Figure 4b). Following doxycycline administration, Rosa-rtTA double Tg mice developed splenomegaly (Figure 4c, d), similar to *St6gal1* single Tg mice. Their skin also exhibited epidermal thinning (Figure 4e, f) and decreased proliferation (Figure 4g, h) comparable to those seen in *Man1a* Tg mice, indicating that the epidermal atrophic phenotype is primarily driven by *Man1a* activity. In contrast, K10/K36 lineage marker staining remained unchanged (Figure 4i, j), consistent with both single Tg lines. Thus, ubiquitous co-induction of *St6gal1* and *Man1a* reproduced both the inflammatory spleen phenotype associated with *St6gal1* expression and the reduced epidermal proliferation and thickness associated with *Man1a* expression.

**Figure 4.**
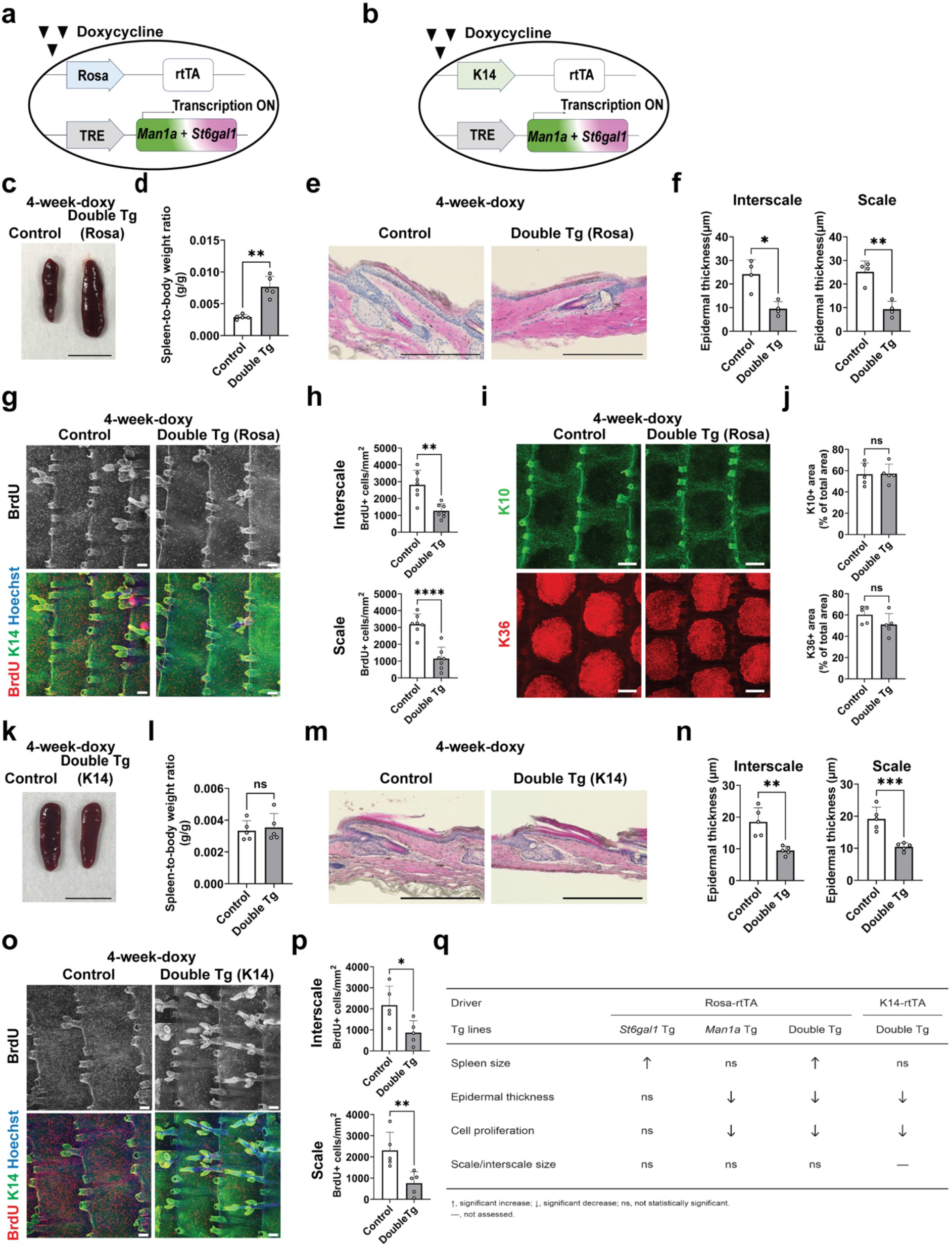
Ubiquitous and epidermis-specific co-expression of *St6gal1* and *Man1a* reduces epidermal thickness and proliferation *in vivo*. (**a, b**) Doxycycline-inducible Tet-On systems used to co-express *St6gal1* and *Man1a* using the ubiquitous Rosa-rtTA driver (a) or the epidermis-specific K14-rtTA driver (b). (**c, d**) Representative spleen picture and spleen/body weight ratio from control and *St6gal1/Man1a* double Tg mice carrying Rosa-rtTA after 4 weeks of doxycycline treatment. Scale bar, 10 mm. (**e, f**) H&E staining of tail skin (e) and quantification of epidermal thickness (f) in control and Rosa-rtTA double Tg mice. Scale bars, 300 µm. (**g, h**) Whole-mount staining of tail epidermis for BrdU (proliferation, red), K14 (epidermal basal cells, green), and Hoechst (blue) (g) and quantification of BrdU-positive epidermal cells (h). Scale bars, 100 µm. (**i, j**) Whole-mount staining for K10 (interscale, green), K36 (scale, red), and Hoechst (blue) (i) and quantification of K10- and K36-positive areas (j). Scale bars, 200 µm. (**k, l**) Representative spleen picture and spleen/body weight ratio from control and *St6gal1/Man1a* double Tg mice carrying K14- rtTA after 4 weeks of doxycycline treatment. Scale bar, 10 mm. (**m, n**) H&E staining of tail skin (m) and quantification of epidermal thickness (n) from control and K14-rtTA double Tg mice. Scale bars, 300 µm. (**o, p**) Whole-mount staining of tail epidermis for BrdU (proliferation, red), K14 (epidermal basal cells, green), and Hoechst (blue) (o) and quantification of BrdU-positive epidermal cells (p). Scale bars, 100 µm. (**q**) Summary of the phenotypes observed across the transgenic mouse models examined in this study. Data are presented as the mean ± SD. Each dot represents one mouse, and measurements from multiple regions of the same mouse are averaged. Comparisons between control and Tg groups were performed using two-tailed unpaired Student’s t-tests. \**P* < 0.05; \*\**P* < 0.01; \*\*\**P* < 0.001; \*\*\*\**P* < 0.0001; ns, not significant.

Since individual and combined expression of *St6gal1* and *Man1a* were induced throughout the whole body when the Rosa-rtTA driver was used, the observed epidermal phenotypes may result from systemic effects of these glycogenes, rather than effects in any particular cell type in the skin. We therefore induced combined *St6gal1* and *Man1a* expression in epidermal basal cells using the K14-rtTA driver (Figure 4b). After 4 weeks of doxycycline administration, the K14-rtTA double Tg mice did not exhibit splenomegaly (Figure 4k, l). Epidermal thickness (Figure 4m, n) and proliferation (Figure 4o, p) were significantly decreased compared with controls, resembling the epidermal phenotypes observed in *Man1a* single Tg and Rosa-rtTA double Tg mice. These findings indicate that both impaired proliferative capacity and epidermal thickness changes may depend on the local function of *St6gal1* and *Man1a* in the epidermis, while splenomegaly may be associated with the systemic effects of *St6gal1* (Figure 4q).

### SNA-based glycoproteomic profiling identifies candidate α2,6-sialylated membrane proteins in keratinocytes co-expressing *St6gal1* and *Man1a*

To identify membrane proteins potentially affected by altered glycosylation, we performed an SNA lectin-based pull-down assay following *St6gal1* and *Man1a* expression (Figure 5a). Primary mouse keratinocytes were transduced with lentiviral vectors encoding *St6gal1* and *Man1a*, and membrane-enriched fractions were subjected to pull-down with SNA, which preferentially recognizes glycoproteins containing terminal α2,6-sialic acid modifications (Figure 5a). SNA-based lectin blotting of the enriched fractions showed a marked increase in signals in *St6gal1*/*Man1a*-co-expressing cells compared with control cells, particularly between approximately 75 and 150 kDa (Figure 5b). SNA-enriched bands in the 75–100-kDa and 100–150-kDa ranges were subsequently analyzed by mass spectrometry.

**Figure 5.**
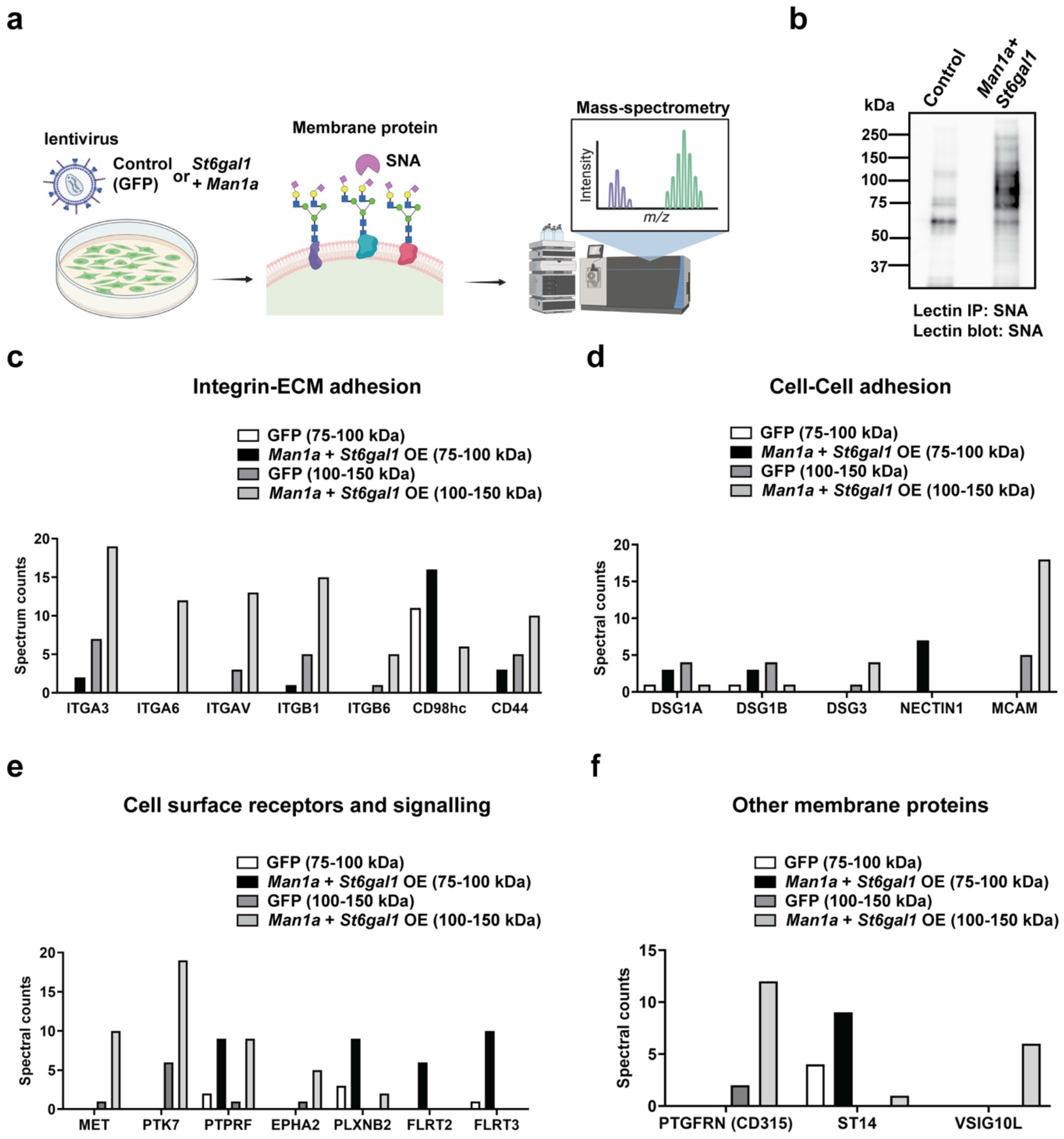
SNA-based glycoproteomic profiling identifies candidate α2,6-sialylated membrane proteins in keratinocytes co-expressing *St6gal1* and *Man1a*. (**a**) Schematic representation of the experimental workflow. Primary mouse keratinocytes were transduced with lentiviral vectors encoding EGFP or *St6gal1* and *Man1a*. Membrane-enriched protein fractions were subjected to SNA-based pull-down, followed by SDS–PAGE and mass spectrometric analysis. (**b**) SNA lectin blotting of membrane-enriched protein fractions from EGFP control and *St6gal1*/Man1a-co-expressing keratinocytes. (**c–f**) Spectral counts of candidate SNA-enriched membrane proteins grouped according to their reported functions: integrin–ECM adhesion (c), cell–cell adhesion (d), cell-surface receptors and signaling-associated proteins (e), and other membrane proteins (f). Candidate proteins were selected according to the criteria described in the Methods.

The SNA-enriched fractions contained several proteins involved in integrin–ECM adhesion, such as integrin subunits (αV, α3, α6, β1, β6), CD98hc, and CD44 (Figure 5c, Supplemental Table S2). Proteins associated with cell–cell adhesion were also detected, including desmosomal junction proteins (DSG1A, DSG1B, DSG3), Nectin-1, and MCAM/CD146 (Figure 5d). Additional candidates included cell-surface receptors and signaling-associated proteins, such as MET, PTK7, PTPRF, EPHA2, PLXNB2, FLRT2, and FLRT3 (Figure 5e), as well as PTGFRN/CD315, ST14, and VSIG10L (Figure 5f). Collectively, these analyses identified candidate SNA-enriched proteins associated with the co-expression of *St6gal1* and *Man1a* in keratinocytes. The enrichment of these candidate α2,6-sialylated proteins, which are involved in cell–ECM adhesion, cell–cell adhesion, and receptor signaling, suggests that these functional groups may be affected by altered glycan modifications. However, further analyses are required to confirm the α2,6-sialylation of individual candidates and determine the functional consequences.

## Discussion

Glycans are important molecules for various biological processes, but their role in aging— particularly in solid tissues, such as skin—remains poorly understood. In this study, we have demonstrated the *in vivo* consequences of increasing the expression of the glycogenes *St6gal1* and *Man1a*, which are upregulated in aged epidermal stem cells (Oinam et al. 2020). Systemic *St6gal1* induction was associated with splenomegaly and an inflammatory transcriptional signature in epidermal basal cells, whereas *Man1a* expression led to local epidermal phenotypes. These findings indicate that distinct N-glycan maturation processes induced by *St6gal1* and *Man1a* differentially influence epidermal homeostasis *in vivo*, although the relationship between these phenotypes and physiological skin aging remains to be established.

Previous studies have reported cell-type-dependent changes in α2,6-sialylation during skin aging: dermal fibroblasts tend to lose α2,6-sialylation under senescent conditions (Sasaki et al. 2017; Itakura et al. 2016), whereas epidermal basal cells exhibit increased α2,6-sialylation with age (Oinam et al. 2020). Such divergent patterns imply that *St6gal1*-mediated glycan remodeling may exert distinct and potentially opposing effects depending on the cell type involved. In our current study, epidermis-specific co-expression of *St6gal1* and *Man1a* using the K14-rtTA driver impaired epidermal homeostasis, supporting the functional relevance of age-associated glycosylation changes in the epidermis. In contrast, the splenomegaly and inflammatory transcriptional changes observed in single *St6gal1* Tg mice likely reflect systemic effects, consistent with the established immunomodulatory functions of *St6gal1* (Appenheimer 2003; Holdbrooks et al. 2020). The absence of splenomegaly in the epidermis-specific double Tg model supports this interpretation, although the cellular origin of the inflammatory phenotypes remains to be determined. Studies with dermis-, immune-, or vascular cell-specific models will be required to further understand the cell-type-specific contributions of glycogenes in skin aging processes.

*St6gal1* and *Man1a* act at different stages of glycan maturation: *Man1a* trims mannose residues to enable subsequent complex N-glycan branching, whereas *St6gal1* primarily catalyzes the addition of terminal α2,6-linked sialic acid to N-glycans (Schjoldager et al. 2020; Moremen et al. 2012). Increased *Man1a* activity may therefore alter the availability of complex N-glycan acceptors for both α2,3- and α2,6-sialyltransferases, which may compete for overlapping substrates (Hwang et al. 2025; Qi et al. 2020). The phenotype of *Man1a* Tg mice potentially reflects broad remodeling of N-glycan maturation, rather than a selective change in α2,6-sialylation. Mannose trimming is also linked to glycoprotein quality control: *Man1a* has been implicated in the targeting of glycoproteins for ER-associated degradation, and increased *Man1a* expression has been associated with activation of the unfolded protein response (Ogen-Shtern et al. 2016; Tu et al. 2017). Altered *Man1a* expression could therefore affect glycoprotein maturation, stability, or trafficking and thereby modify the cell-surface localization or function of proteins involved in epidermal homeostasis. In addition, although *St6gal1* preferentially modifies N-glycans, it could potentially affect other glycan structures, including O-glycans (Cherian et al. 2015). Thus, the combined phenotype of double Tg mice may arise from complex interactions among mannose trimming, glycoprotein quality control, and terminal sialylation pathways, rather than from a simple linear cascade. Detailed structural glycomic and mechanistic analyses will be required to determine how *St6gal1* and/or *Man1a* expression affects glycan structure and glycoprotein processing in the regulation of skin function and the aging process.

SNA-based enrichment followed by mass spectrometry identified candidate α2,6-sialylated membrane proteins in primary keratinocytes co-expressing *St6gal1* and *Man1a*. SNA-enriched membrane proteins were associated with cell–ECM adhesion, cell–cell junctions, or receptor signaling. These included several integrin subunits (αV, α3, α6, β1, β6), together with the integrin-associated protein CD98hc and the adhesion receptor CD44. Integrins are central regulators of epidermal stem cells (Kaur & Li 2000), and their N-glycan composition modulates integrin conformation, ligand binding affinity, clustering, and downstream signal propagation (Kaburagi et al. 2017; Pan & Song 2010; Cai et al. 2017). CD98hc is reported to be co-localized with β1 integrin, and their interaction regulates basal keratinocyte adhesion (Lemaître et al. 2011). Meanwhile, hyaluronan–CD44 signaling has been implicated in barrier function in aged skin (Bourguignon et al. 2013). Thus, altered α2,6-sialylation of integrins may influence the interactions of keratinocytes with the extracellular environment.

The SNA-enriched fractions contained junction-associated proteins as well as cell-surface receptors. Examples include MET, which contributes to keratinocyte migration and wound repair; its N-glycosylation has been reported to affect receptor stability and signaling output (Qian et al. 2009; Li et al. 2013; Saitou et al. 2022). MCAM (MUC18/CD146), an immunoglobulin superfamily adhesion molecule, is typically low or absent in the healthy epidermis but is induced in inflammatory skin diseases (Weninger et al. 2000). Glycosylation has also been implicated in desmosome stability and keratinocyte cell–cell adhesion, while low DSG3 expression has been associated with keratinocytes displaying stem/progenitor-like properties (Wan et al. 2003; Hu et al. 2006; Jin & Chung 2018). Collectively, these findings identify multiple membrane proteins as candidate mediators of glycan-dependent epidermal phenotypes, although further validation of individual candidates is required to establish their direct α2,6-sialylation and related functional consequences.

### Limitations of this study

Several limitations should be considered. First, transgene induction in young adult mice may not fully reproduce the expression levels of glycogenes or glycome changes associated with physiological skin aging. Thus, our findings do not establish whether endogenous upregulation of *St6gal1* or *Man1a* is required for age-associated epidermal changes. Second, glycoproteomic profiling was performed using cultured primary keratinocytes, because the limited protein yield from FACS-isolated epidermal stem cells precluded lectin-based immunoprecipitation and mass spectrometry analyses. Third, current genetic approaches do not allow the selective modification of defined glycan structures on individual membrane proteins *in vivo*. Future studies incorporating glycosylation-site mutants or targeted chemical glycan modifications may help resolve protein-specific contributions. Finally, glycosylation patterns and their function may differ between mice and humans. Therefore, species-specific profiling of glycan structures and validation in human keratinocytes or skin tissue will be essential to establish the translational relevance of our findings.

## Materials and methods

### Animals and ethical approval

All mice were maintained under specific pathogen-free conditions at the Center for Animal Resources and Development (CARD), Kumamoto University, or the Laboratory of Embryonic and Genetic Engineering, Kyushu University. Animals were housed under controlled environmental conditions with a 12-h light/12-h dark cycle and had free access to standard chow and water. C57BL/6J mice used for the maintenance of the transgenic lines were obtained from Charles River Laboratories Japan, Inc. (Kanagawa, Japan) or Japan SLC, Inc. (Shizuoka, Japan), as appropriate. Both male and female mice were included in the experiments and were randomly assigned to experimental groups. All procedures involving animals were performed in accordance with the institutional guidelines of Kumamoto University and Kyushu University and were approved by their respective animal experimentation committees.

### Generation and doxycycline treatment of inducible *St6gal1* and *Man1a* Tg mice

Doxycycline-inducible *St6gal1* and *Man1a* Tg mice were generated using a Tet-On system. Mouse *St6gal1* and *Man1a* coding sequences carrying an N-terminal Flag or Myc tag, respectively, were cloned downstream of the tetracycline-responsive element (TRE) in the pTRE2 vector (Clontech). The pTRE–Flag-*St6gal1* and pTRE–Myc-*Man1a* expression cassettes were isolated from the vector backbone and purified for pronuclear microinjection into C57BL/6J zygotes. Transgenic mice were generated at the CARD, Kumamoto University.

Genomic integration of each transgene was assessed by PCR using primers specific for the corresponding pTRE–transgene cassette: pTRE–Flag-*St6gal1*, forward 5′-TCGCCTGGAGACGCCAT-3′ and reverse 5′-TTCTCCTGGCTCTTCGGCATC-3′; pTRE–Myc-*Man1a*, forward 5′-AAGTGAAAGTCGAGCTCGGTA-3′ and reverse 5′-GAAGCAGAGCGTGATGAAGG-3′. The founder lines used in this study were selected on the basis of stable germline transmission and reproducible doxycycline-dependent transgene induction.

The pTRE–Flag-*St6gal1* and pTRE–Myc-*Man1a* Tg mice were crossed with Rosa-rtTA mice (JAX #006965) to induce ubiquitous transgene expression or with K14-rtTA mice (JAX #008099) to induce expression in epidermal basal cells. The K14-rtTA line was backcrossed to C57BL/6J mice for at least eight generations before experimental use. Mice carrying both pTRE–Flag-*St6gal1* and pTRE–Myc-*Man1a* transgenes were used for co-expression and are referred to as double Tg mice. Transgene expression was induced beginning at 2–3 months of age by administering custom-formulated chow containing doxycycline hyclate at 1 g/kg (D9891; Sigma-Aldrich), prepared by Oriental Yeast Co., Ltd., for the indicated durations. Control mice received the same doxycycline-containing chow for the same duration as the respective experimental groups.

### Flag/Myc immunoprecipitation and immunoblotting using epidermal cells

Epidermal cells were isolated as previously described with minor modifications (Oinam et al. 2020; Fan et al. 2026), with back skin being used instead of tail skin. After removal of the subcutaneous tissue and fat, the skin was rinsed with 1× PBS and incubated in 0.25% trypsin/EDTA overnight at 4 °C, followed by incubation at 37 °C for 30 min. Single-cell suspensions were prepared by gentle scraping of the epidermis. The collected cells were washed twice with 1× PBS and then stored at −80 °C until protein extraction and immunoprecipitation.

Epidermal cell pellets (1 × 10^6^ cells) were thawed on ice and lysed in 200–300 µL of Pierce IP Lysis Buffer (Thermo Fisher Scientific) supplemented with a protease inhibitor cocktail (Sigma-Aldrich) and Phosphatase Inhibitor Cocktail Solution I (FUJIFILM Wako Pure Chemical Corporation). After vortexing, the lysates were incubated on ice for 10 min and subjected to approximately 10 cycles of sonication using an ultrasound homogenizer Q125 (Wakanebee Tech Co., Ltd.), with 10-s pulses separated by 59-s intervals, while maintained on ice. Protein concentrations were measured using the Pierce BCA Protein Assay Kit (Thermo Fisher Scientific).

For each control–Tg comparison in immunoprecipitation, lysates were normalized to the same total protein input, and equal amounts of total protein were used for all samples within a given experiment. An aliquot corresponding to 10% of each lysate was retained as the input. The remaining lysates were incubated with 30–40 µL of anti-FLAG M2 affinity gel (Sigma-Aldrich, A2220) or anti-c-Myc agarose conjugate (Nacalai Tesque, 04145-55) for 2 h at 4 °C with rotation. Following incubation, the beads were washed four times with Pierce IP Lysis Buffer and collected by centrifugation at 4 °C, 5,000 rpm for 1 min. Bound proteins were eluted by adding 30 µL of 1.5× SDS buffer and heating at 95 °C for 5 min.

Eluted proteins and input samples were separated by SDS–PAGE using a 10% gel and transferred to a PVDF membrane using a semi-dry transfer system (Bio-Rad PowerPac HC). The membranes were blocked with 3% Amersham ECL Blocking Agent for 1 h at room temperature and incubated for 1 h at room temperature with rabbit anti-Flag antibody (Cell Signaling Technology, 14793, 1:2,000) or rabbit anti-Myc antibody (Cell Signaling Technology, 2272, 1:2,000). After three 5-min washes with TBS containing 0.1% Tween 20 (TBS-T), the membranes were incubated with HRP-conjugated goat anti-rabbit IgG antibody (Bio-Rad, 1606515, 1:5,000) for 1 h at room temperature. Following three additional washes, chemiluminescent signals were developed using ECL Western Blotting Detection Reagents (Cytiva, RPN418) and captured using an ImageQuant LAS 4000 mini system (GE Healthcare).

### FACS isolation and lectin blotting of epidermal basal cells

Epidermal basal cells were isolated from tail skin by fluorescence-activated cell sorting (FACS), as previously described (Oinam et al. 2020; Fan et al. 2026). Briefly, epidermal cell suspensions were prepared and stained on ice for 30 min with biotin-conjugated anti-CD34 antibody (eBioscience, 13-0341-85, 1:50), APC-conjugated streptavidin (BD Biosciences, 554067, 1:100), BV510-conjugated anti-integrin α6 antibody (BD Biosciences, 563271, 1:200), and BV421-conjugated anti-Sca-1 antibody (BD Biosciences, 562729, 1:100). Dead cells were excluded using 7-AAD (BD Biosciences). Viable integrin α6^high^/CD34^−^/Sca-1^+^ epidermal basal cells (300,000 cells per tube) were isolated into 5% BCS/PBS solution using a FACSAria flow cytometer (BD Biosciences). After centrifugation at 12,000 rpm for 5 min at 4 °C, collected cell pellets were washed three times with ice-cold 1× PBS and stored at −80 °C. Flow cytometry data were analyzed using FlowJo software (BD Biosciences).

Membrane-enriched hydrophobic and hydrophilic fractions were prepared from the sorted epidermal basal cells using the CelLytic MEM Protein Extraction Kit (Sigma-Aldrich, CE0050) according to the manufacturer’s instructions. Only the membrane-enriched hydrophobic fractions were used for lectin blotting.

Recombinant rHeltuba (a mannose binder) was prepared as previously described (Tateno et al. 2011), and SNA (EBL, Funakoshi L-1300; α2,6-sialic acid binder) was used for the lectin blot. The lectins were conjugated to horseradish peroxidase (HRP) using an HRP Labeling Kit (Dojindo Molecular Technologies) according to the manufacturer’s instructions. HRP-conjugated lectins were stored at 0.75 µg/µL in 50% glycerol at −20 °C until use.

Equal amounts of protein (2 µg) from the membrane-enriched hydrophobic fractions were separated by SDS–PAGE using a Perfect NT Gel System (DRC, NTH-676HP) and transferred to PVDF membranes using a wet-transfer system (Bio-Rad). The membrane was blocked for 1 h at room temperature with Carbo-Free Blocking Solution (Vector Laboratories) containing 0.1% Tween 20. The membrane was then incubated for 1 h at room temperature with HRP-conjugated rHeltuba or SNA at a final concentration of 0.75 µg/mL in 1× TBS-T. After three washes with 1× TBS-T, signals were developed using ECL Western Blotting Detection Reagents (Cytiva, RPN418) and captured using an ImageQuant LAS 4000 mini system (GE Healthcare).

### Body and spleen weight measurements

At the experimental endpoint, mice were euthanized, and their body and spleen weights were recorded. Relative spleen weight was calculated as the spleen weight normalized to the body weight of each mouse.

### Histological and whole-mount analyses of tail skin

Histological and whole-mount analyses of tail skin were performed as previously described (Sada et al. 2016; Fan et al. 2026). For histological analysis, tail skin was embedded in OCT compound (Tissue-Tek; Sakura Finetek), sectioned at 10 µm, fixed with 4% paraformaldehyde for 10 min at room temperature, and stained with hematoxylin and eosin (H&E). Images were acquired using an EVOS M5000 Imaging System (Thermo Fisher Scientific). Epidermal thickness was measured in H&E-stained sections using Fiji/ImageJ (National Institutes of Health). For each mouse, 6–16 epidermal structures were analyzed, as previously described (Changarathil et al. 2019). Measurements from multiple epidermal structures were averaged to obtain a value per mouse for statistical analysis.

For proliferation analysis, BrdU (Sigma-Aldrich, B5002) was administered in drinking water at 0.8 mg/mL for 2 days before euthanasia. Tail epidermal whole mounts were stained with rat anti-BrdU (Abcam, ab6326, 1:300), rabbit anti-K14 (BioLegend, 905304, 1:1,000), mouse anti-K10 (Abcam, ab9026, 1:500), or rabbit anti-K36 (Proteintech, 14309-1-AP, 1:300) antibodies. Samples were imaged by Z-stack acquisition using a Nikon A1 HD25 or Zeiss LSM900 confocal microscope.

Whole-mount images were quantified using Fiji/ImageJ. For BrdU analysis, 6–12 fields containing 8–16 scale and interscale regions were measured per mouse, and BrdU-positive cells were counted manually. For K10 and K36 analyses, 4–10 fields containing a total of 12– 20 epidermal units were analyzed per mouse. Each epidermal unit comprises one scale and one adjacent interscale region. Measurements from multiple fields or epidermal regions were averaged to obtain a value per mouse for statistical analysis.

### FACS isolation of epidermal basal cells and RNA sequencing

Epidermal basal cells were isolated from tail skin by FACS, as described above. Viable integrin α6^high^/CD34^−^/Sca-1^+^ cells isolated from control and pTRE–Flag-*St6gal1*/Rosa-rtTA mice (n = 3 mice per group) were sorted directly into TRIzol LS Reagent (Ambion).

RNA extraction, library preparation, sequencing, and primary data processing were performed by Tsukuba i-Laboratory LLP (University of Tsukuba), as previously described (Fan et al. 2026). Briefly, RNA quantity and quality were assessed using a NanoDrop spectrophotometer and an Agilent 2100 Bioanalyzer with the RNA 6000 Pico Kit. Libraries were prepared using the SMARTer Stranded Total RNA-Seq Kit v2—Pico Input Mammalian (Takara Bio) and sequenced on an Illumina NextSeq 500 platform. Sequencing reads were mapped to the annotated mouse reference genome using CLC Genomics Workbench version 20.0.3 (Qiagen), and read counts were normalized across samples by quantile normalization. Fold changes were calculated from log_2_-transformed normalized expression values, and genes exhibiting an absolute fold change of ≥2 were used for downstream analyses.

PCA was performed using CLC Genomics Workbench. Gene Ontology enrichment analysis was conducted using Metascape (https://metascape.org) with genes exhibiting an absolute fold change of ≥2. Hierarchical clustering and heatmap visualization were performed using Morpheus (Broad Institute) (https://software.broadinstitute.org/morpheus). Gene set enrichment analysis was conducted using GSEA version 4.4.0 (Subramanian et al. 2005).

### Primary mouse keratinocyte culture and lentiviral transduction

Primary keratinocytes were isolated from the neonatal skin of C57BL/6J mice and cultured as previously described (Lichti et al. 2008; Oinam et al. 2020). Lentiviral vectors encoding mouse *St6gal1*, *Man1a*, or EGFP were generated using the CSII-CMV-MCS-IRES2-Bsd vector and prepared as previously described (Oinam et al. 2020). Primary keratinocytes were seeded on collagen IV-coated culture plates and cotransduced with lentiviruses encoding *St6gal1* and *Man1a* in the presence of 4 µg/mL polybrene for 16 h. Control cells were transduced with an EGFP-expressing lentivirus. After replacement with fresh medium, transduced cells were selected with 1 µg/mL blasticidin. Membrane-enriched protein fractions prepared from the transduced keratinocytes were subsequently used for the SNA-based pull-down assay and mass spectrometry analysis.

### SNA lectin-based pull-down assay and mass spectrometry analysis

SNA was biotinylated using the Biotin Labeling Kit-NH₂ (Dojindo, LK03) according to the manufacturer’s instructions, and immobilized on Dynabeads M-280 Streptavidin (Invitrogen). For each sample, 10 µL of beads were incubated with biotinylated SNA for 2 h at 4 °C with shaking. The SNA-conjugated beads were collected using a DynaMag-2 magnet (Invitrogen) and washed with 1× PBS containing 1% Triton X-100. Equal amounts of membrane-enriched protein from primary keratinocytes cotransduced with lentiviruses encoding *St6gal1* and *Man1a* or transduced with an EGFP-expressing control vector were incubated with the SNA-conjugated beads overnight at 4 °C with shaking. The beads were collected and washed four times with 1% PBS-T. Bound proteins were released by heating the beads in 0.2% SDS at 95 °C for 5 min. Residual biotinylated SNA in the eluates was removed by incubation with fresh Dynabeads M-280 Streptavidin for 1 h at room temperature.

Equal amounts of the SNA-enriched proteins were separated by SDS–PAGE using a Perfect NT Gel System (DRC, NTH-676HP) and visualized using a mass spectrometry-compatible silver-staining kit (FUJIFILM Wako Pure Chemical Corporation). Gel regions corresponding to 75–100 kDa and 100–150 kDa were excised separately from the EGFP control and *St6gal1*/*Man1a*-co-expression samples. Protein identification by nanoLC–MS/MS was performed by Kazusa Genome Technologies Inc. (Kisarazu, Japan).

The excised gel pieces were destained, dehydrated with acetonitrile, and digested overnight at 37 °C with Trypsin/Lys-C Mix (Promega). Peptides were extracted with 5% formic acid in 50% acetonitrile, desalted using MonoSpin C18 columns (GL Sciences), and analyzed using an UltiMate 3000 RSLCnano LC system coupled to a Q Exactive HF-X mass spectrometer (Thermo Fisher Scientific). Mass spectra were acquired in positive-ion mode using data-dependent acquisition with higher-energy collisional dissociation.

Protein identification was performed using PEAKS Studio X+ (Bioinformatics Solutions Inc.) against the reviewed mouse UniProtKB/Swiss-Prot database (proteome ID UP000000589; 21,949 entries; downloaded March 7, 2023). Database searches were conducted with precursor and fragment mass tolerances of 10 ppm and 0.015 Da, respectively, allowing up to two missed tryptic cleavages. Methionine oxidation was included as a variable modification.

The exported protein-identification results were filtered in Microsoft Excel. Candidate proteins were required to be identified by at least four unique peptides and to meet one of the following spectral-count criteria in at least one of the two gel regions: (1) a *St6gal1*/*Man1a* co-expression to EGFP control spectral-count ratio of ≥2; or (2) when no spectra were detected in the corresponding control sample, at least five spectra in the *St6gal1*/*Man1a*-co-expression sample. The resulting proteins were manually reviewed for reported plasma-membrane localization, and proteins classified as plasma-membrane proteins were retained as final candidates.

### Statistical analysis

Statistical analyses were applied to spleen-to-body-weight ratios, epidermal thickness measurements, and quantitative analyses of whole-mount staining. Unless otherwise indicated, at least three mice were analyzed per group. Measurements obtained from multiple images or epidermal regions from the same mouse were averaged, and each mouse was treated as one biological replicate and one statistical unit. Data are presented as the mean ± standard deviation (SD), with individual data points representing individual mice.

Comparisons between control and transgenic groups were performed using two-tailed unpaired Student’s t-tests in GraphPad Prism 9 (GraphPad Software). A *P* value < 0.05 was considered statistically significant. Significance is indicated as follows: \**P* < 0.05; \*\**P* < 0.01; \*\*\**P* < 0.001; \*\*\*\**P* < 0.0001; ns, not significant.

## Data Availability

The RNA sequencing datasets have been deposited in the National Center for Biotechnology Information-Gene Expression Omnibus (GEO) under accession number GSE313192 (https://www.ncbi.nlm.nih.gov/geo/query/acc.cgi?acc=GSE313192).

All raw mass spectrometry data files have been deposited in the Mass Spectrometry Interactive Virtual Environment (MassIVE), hosted by the Center for Computational Mass Spectrometry at the University of California, San Diego, and are publicly available under accession number MSV000103629 (https://massive.ucsd.edu/ProteoSAFe/dataset.jsp?accession=MSV000103629).

## Acknowledgments

We thank the International Core Facility of Advanced Life Science at Kumamoto University and the Research Promotion Unit at Kyushu University for their kind support with the core facility. We also thank the CARD at Kumamoto University and the Laboratory of Embryonic and Genetic Engineering at Kyushu University for their support in the care and management of laboratory animals. We thank K. Araki and N. Takeda (Kumamoto University) for their valuable support in generating transgenic mouse lines. We thank G. Sashida, T. Keida (Kumamoto University), N. Imai, and M. Huang (Kyushu University) for their technical assistance. We thank S. Nishihara, A. Togayachi, and K. Angata (Soka University) for insightful discussions about glycogenes and glycan biology. We also thank G. Sheng, M. Ogawa, Y. Arima, and K. Miharada (Kumamoto University) for helpful discussions and critical feedback on this work. We also thank A. Yesbolatova (Kyushu University) for the critical reading of this manuscript.

This work was supported by AMED-PRIME, Japan Agency for Medical Research and Development (AMED; JP25gm6710011 and JP21gm6110016), JSPS KAKENHI, Grant-in-Aid for Scientific Research (B) (JP24K02035), and Grant-in-Aid for Challenging Research (Exploratory) (JP20K21431) (all to A.S.). Additional funding to A.S. was provided by the Takeda Science Foundation, the Mitsubishi Foundation, KOSE Cosmetology Research Foundation, the Senri Life Science Foundation, Astellas Foundation for Research on Metabolic Disorders, Tokyo Biochemical Research Foundation, the Sumitomo Foundation, the Nakatomi Foundation, and Hoyu Science Foundation. T.B.S. received scholarship support from the Ministry of Education, Culture, Sports, Science and Technology of Japan (MEXT), and A.G.K.H. was supported by the Egypt–Japan Education Partnership (EJEP) Scholarship.

This work was also supported in part by the MEXT Cooperative Research Project Program, Medical Research Center Initiative for High Depth Omics, and CURE (JPMXP1323015486) to the Medical Institute of Bioregulation (MIB), Kyushu University. The infrastructure of Omics Science Center Secure Information Analysis System (OASIS; https://sis.bioreg.kyushu-u.ac.jp/), Medical Institute of Bioregulation at Kyushu University, provided part of the computational resources.

## Author Contributions

A.S., H.T., and H.Y. conceived and designed the study. T.B.S., N.T.K.N., E.R., M.I., A.G.K.H., and L.O. performed experiments. T.B.S., N.T.K.N., and A.S. analyzed and interpreted the data. T.B.S. and N.T.K.N. validated the experimental results and curated the data. N.T.K.N. and A.S. designed and validated the transgenic mouse lines. A.S. developed and provided the methodologies and resources for the mouse and skin phenotype analyses. H.T. provided lectin-related materials and protocols and contributed to the development of the lectin-based methodology. A.S. supervised the study, administered the project, and acquired funding. T.B.S., N.T.K.N., and A.S. prepared the figures. T.B.S. and N.T.K.N. wrote the original draft. A.S., M.I., H.T., and H.Y. reviewed and edited the manuscript. All authors read and approved the final manuscript.

## Declaration of Generative AI and AI-assisted Technologies in the Writing Process

During the preparation of this manuscript, the authors used ChatGPT to improve clarity and readability. The authors subsequently reviewed and revised the manuscript and take full responsibility for its final content.

## Supplementary information

**Supplemental Table S1.** List of differentially expressed genes identified by RNA-seq analysis of epidermal basal cells from *St6gal1* Tg and control mice (related to Figure 3).

**Supplemental Table S2.** List of identified peptides from EGFP- and *St6gal1*/*Man1a*-expressing mouse keratinocytes (related to Figure 5c–f).

## References

Abdayem R, Formanek F, Minondo AM, Potter A & Haftek M (2016) Cell surface glycans in the human stratum corneum: distribution and depth-related changes. Experimental Dermatology 25, 865–871.

Ankenbauer KE, Rao TC, Mattheyses AL & Bellis SL (2023) Sialylation of EGFR by ST6GAL1 induces receptor activation and modulates trafficking dynamics. J Biol Chem 299, 105217.

Appenheimer MM, Huang R-Y, Chandrasekaran EV, Dalziel M, Hu YP, Soloway PD, Wuensch SA, Matta KL & Lau JTY (2003) Biologic contribution of P1 promoter– mediated expression of ST6Gal I sialyltransferase. Glycobiology 13, 591–600.

Bourguignon LYW, Wong G, Xia W, Man M-Q, Holleran WM & Elias PM (2013) Selective Matrix (Hyaluronan) Interaction with CD44 and RhoGTPase Signaling Promotes Keratinocyte Functions and Overcomes Age-related Epidermal Dysfunction. J Dermatol Sci 72, 32–44.

Broussard AC & Boyce M (2019) Life is sweet: the cell biology of glycoconjugates. Mol Biol Cell 30, 525–529.

Cai X, Thinn AMM, Wang Z, Shan H & Zhu J (2017) The importance of N-glycosylation on β3 integrin ligand binding and conformational regulation. Scientific Reports 7, 4656.

Catera M, Borelli V, Malagolini N, Chiricolo M, Venturi G, Reis CA, Osorio H, Abruzzo PM, Capri M, Monti D, Ostan R, Franceschi C & Dall’Olio F (2016) Identification of novel plasma glycosylation-associated markers of aging. Oncotarget 7, 7455–7468.

Changarathil G, Ramirez K, Isoda H, Sada A & Yanagisawa H (2019) Wild-type and SAMP8 mice show age-dependent changes in distinct stem cell compartments of the interfollicular epidermis. PLoS One 14, e0215908.

Cherian RM, Jin C, Liu J, Karlsson NG & Holgersson J (2015) A Panel of Recombinant Mucins Carrying a Repertoire of Sialylated O-Glycans Based on Different Core Chains for Studies of Glycan Binding Proteins. Biomolecules 5, 1810–1831.

Dabelsteen S, Pallesen EMH, Marinova IN, Nielsen MI, Adamopoulou M, Rømer TB, Levann A, Andersen MM, Ye Z, Thein D, Bennett EP, Büll C, Moons SJ, Boltje T, Clausen H, Vakhrushev SY, Bagdonaite I & Wandall HH (2020) Essential Functions of Glycans in Human Epithelia Dissected by a CRISPR-Cas9-Engineered Human Organotypic Skin Model. Dev Cell 54, 669–684.e7.

Donovan M, Duke R, Simonetti L, Cavusoglu N, Rudd PM & Bernard D (2023) N-Glycans Are Stratum Corneum Biomarkers of Aging Skin. Journal of Investigative Dermatology 143, 492–494.e10.

Fan W, Ishikawa M, Raja E, Hegazy AM, Date H, Ngo YX, Sato Y, Yamagata K, Yanagisawa H & Sada A (2026) Integrin-Binding Matricellular Protein Fibulin-5 Maintains Epidermal Stem Cell Heterogeneity During Skin Aging. Aging Cell 25, e70483.

Giangreco A, Qin M, Pintar JE & Watt FM (2008) Epidermal stem cells are retained in vivo throughout skin aging. Aging Cell 7, 250–259.

Gilchrest BA (1983) In Vitro Assessment of Keratinocyte Aging. Journal of Investigative Dermatology 81, S184–S189.

Gomez C, Chua W, Miremadi A, Quist S, Headon DJ & Watt FM (2013) The Interfollicular Epidermis of Adult Mouse Tail Comprises Two Distinct Cell Lineages that Are Differentially Regulated by Wnt, Edaradd, and Lrig1. Stem Cell Reports 1, 19–27.

Gu J & Taniguchi N (2004) Regulation of integrin functions by N-glycans. Glycoconjugate Journal 21, 9–15.

Hanisch F, Weidemann W, Großmann M, Joshi PR, Holzhausen H-J, Stoltenburg G, Weis J, Zierz S & Horstkorte R (2013) Sialylation and Muscle Performance: Sialic Acid Is a Marker of Muscle Ageing. PLoS One 8, e80520.

Holdbrooks AT, Ankenbauer KE, Hwang J & Bellis SL (2020) Regulation of inflammatory signaling by the ST6Gal-I sialyltransferase. PLoS One 15, e0241850.

Holdbrooks AT, Britain CM & Bellis SL (2018) ST6Gal-I sialyltransferase promotes tumor necrosis factor (TNF)-mediated cancer cell survival via sialylation of the TNF receptor 1 (TNFR1) death receptor. J Biol Chem 293, 1610–1622.

Hu P, Berkowitz P, Madden VJ & Rubenstein DS (2006) Stabilization of Plakoglobin and Enhanced Keratinocyte Cell-Cell Adhesion by Intracellular O-Glycosylation. Journal of Biological Chemistry 281, 12786–12791.

Hwang J, Rao TC, Tao J, Sha B, Narimatsu Y, Clausen H, Mattheyses AL & Bellis SL (2025) Apoptotic signaling by TNFR1 is inhibited by the α2-6 sialylation, but not α2-3 sialylation, of the TNFR1 *N*-glycans. Journal of Biological Chemistry 301, 108043.

Itakura Y, Sasaki N, Kami D, Gojo S, Umezawa A & Toyoda M (2016) N- and O-glycan cell surface protein modifications associated with cellular senescence and human aging. Cell Biosci 6, 14.

Jin S-P & Chung JH (2018) Inhibition of N-glycosylation by tunicamycin attenuates cell–cell adhesion via impaired desmosome formation in normal human epidermal keratinocytes. Biosci Rep 38, BSR20171641.

Kaburagi T, Kizuka Y, Kitazume S & Taniguchi N (2017) The Inhibitory Role of α2,6-Sialylation in Adipogenesis*. Journal of Biological Chemistry 292, 2278–2286.

Kaur P & Li A (2000) Adhesive Properties of Human Basal Epidermal Cells: An Analysis of Keratinocyte Stem Cells, Transit Amplifying Cells, and Postmitotic Differentiating Cells. Journal of Investigative Dermatology 114, 413–420.

Keyes BE, Liu S, Asare A, Naik S, Levorse J, Polak L, Lu CP, Nikolova M, Pasoli HA & Fuchs E (2016) Impaired Epidermal to Dendritic T-Cell Signaling Slows Wound Repair in Aged Skin. Cell 167, 1323–1338.e14.

Kornfeld R & Kornfeld S (1985) Assembly of Asparagine-Linked Oligosaccharides. Annual Review of Biochemistry 54, 631–664.

Krick S, Helton ES, Easter M, Bollenbecker S, Denson R, Zaharias R, Cochran P, Vang S, Harris E, Wells JM & Barnes JW (2021) ST6GAL1 and α2-6 Sialylation Regulates IL-6 Expression and Secretion in Chronic Obstructive Pulmonary Disease. Front Immunol 12, 693149.

Legler K, Rosprim R, Karius T, Eylmann K, Rossberg M, Wirtz RM, Müller V, Witzel I, Schmalfeldt B, Milde-Langosch K & Oliveira-Ferrer L (2018) Reduced mannosidase MAN1A1 expression leads to aberrant N-glycosylation and impaired survival in breast cancer. British Journal of Cancer 118, 847–856.

Lemaître G, Stella A, Feteira J, Baldeschi C, Vaigot P, Martin MT, Monsarrat B & Waksman G (2011) CD98hc (SLC3A2) is a key regulator of keratinocyte adhesion. Journal of Dermatological Science 61, 169–179.

Li J-F, Duan H-F, Wu C-T, Zhang D-J, Deng Y, Yin H-L, Han B, Gong H-C, Wang H-W & Wang Y-L (2013) HGF Accelerates Wound Healing by Promoting the Dedifferentiation of Epidermal Cells through β 1-Integrin/ILK Pathway. Biomed Res Int 2013, 470418.

Lichti U, Anders J & Yuspa SH (2008) Isolation and short term culture of primary keratinocytes, hair follicle populations, and dermal cells from newborn mice and keratinocytes from adult mice, for in vitro analysis and for grafting to immunodeficient mice. Nat Protoc 3, 799–810.

Liu J, Dong X, Xie R, Tang Y, Thomas AM, Li S, Liu S, Yu M & Qin H (2024) N-linked α2,6-sialylation of integrin β1 by the sialyltransferase ST6Gal1 promotes cell proliferation and stemness in gestational trophoblastic disease. Placenta 149, 18–28.

López-Otín C, Blasco MA, Partridge L, Serrano M & Kroemer G (2023) Hallmarks of aging: An expanding universe. Cell 186, 243–278.

Möginger U, Grunewald S, Hennig R, Kuo C-W, Schirmeister F, Voth H, Rapp E, Khoo K- H, Seeberger PH, Simon JC & Kolarich D (2018) Alterations of the Human Skin N- and O-Glycome in Basal Cell Carcinoma and Squamous Cell Carcinoma. Front Oncol 8, 70.

Moremen KW, Tiemeyer M & Nairn AV (2012) Vertebrate protein glycosylation: diversity, synthesis and function. Nat Rev Mol Cell Biol 13, 448–462.

Nasirikenari M, Chandrasekaran EV, Matta KL, Segal BH, Bogner PN, Lugade AA, Thanavala Y, Lee JJ & Lau JTY (2010) Altered eosinophil profile in mice with ST6Gal-1 deficiency: an additional role for ST6Gal-1 generated by the P1 promoter in regulating allergic inflammation. J Leukoc Biol 87, 457–466.

Nishihara S (2018) Glycans in stem cell regulation: from Drosophila tissue stem cells to mammalian pluripotent stem cells. FEBS Letters 592, 3773–3790.

Ogen-Shtern N, Avezov E, Shenkman M, Benyair R & Lederkremer GZ (2016) Mannosidase IA is in Quality Control Vesicles and Participates in Glycoprotein Targeting to ERAD. Journal of Molecular Biology 428, 3194–3205.

Oinam L, Changarathil G, Raja E, Ngo YX, Tateno H, Sada A & Yanagisawa H (2020) Glycome profiling by lectin microarray reveals dynamic glycan alterations during epidermal stem cell aging. Aging Cell 19, e13190.

Pan D & Song Y (2010) Role of Altered Sialylation of the I-Like Domain of β1 Integrin in the Binding of Fibronectin to β1 Integrin: Thermodynamics and Conformational Analyses. Biophys J 99, 208–217.

Qi F, Isaji T, Duan C, Yang J, Wang Y, Fukuda T & Gu J (2020) ST3GAL3, ST3GAL4, and ST3GAL6 differ in their regulation of biological functions via the specificities for the α2,3-sialylation of target proteins. The FASEB Journal 34, 881–897.

Qian J, Zhu C-H, Tang S, Shen A-J, Ai J, Li J, Geng M-Y & Ding J (2009) alpha2,6-hyposialylation of c-Met abolishes cell motility of ST6Gal-I-knockdown HCT116 cells. Acta Pharmacol Sin 30, 1039–1045.

Raja E, Changarathil G, Oinam L, Tsunezumi J, Ngo YX, Ishii R, Sasaki T, Imanaka-Yoshida K, Yanagisawa H & Sada A (2022) The extracellular matrix fibulin 7 maintains epidermal stem cell heterogeneity during skin aging. The EMBO Reports 23, EMBR202255478.

Rao TC, Beggs RR, Ankenbauer KE, Hwang J, Ma VP-Y, Salaita K, Bellis SL & Mattheyses AL (2022) ST6Gal-I–mediated sialylation of the epidermal growth factor receptor modulates cell mechanics and enhances invasion. J Biol Chem 298, 101726.

Reily C, Stewart TJ, Renfrow MB & Novak J (2019) Glycosylation in health and disease. Nat Rev Nephrol 15, 346–366.

Sada A, Jacob F, Leung E, Wang S, White BS, Shalloway D & Tumbar T (2016) Defining the cellular lineage hierarchy in the inter-follicular epidermis of adult skin. Nat Cell Biol 18, 619–631.

Saitou A, Hasegawa Y, Fujitani N, Ariki S, Uehara Y, Hashimoto U, Saito A, Kuronuma K, Matsumoto K, Chiba H & Takahashi M (2022) N-glycosylation regulates MET processing and signaling. Cancer Sci 113, 1292–1304.

Sasaki N, Itakura Y & Toyoda M (2017) Sialylation regulates myofibroblast differentiation of human skin fibroblasts. Stem Cell Res Ther 8, 81.

Schjoldager KT, Narimatsu Y, Joshi HJ & Clausen H (2020) Global view of human protein glycosylation pathways and functions. Nat Rev Mol Cell Biol 21, 729–749.

Shen L, Lin J-M, Lin J & Wu W (2025) Glycosylation in Dermatology: Unveiling the Sugar Coating of Skin Disease. Experimental Dermatology 34, e70098.

Shibuya N, Goldstein IJ, Broekaert WF, Nsimba-Lubaki M, Peeters B & Peumans WJ (1987) The elderberry (Sambucus nigra L.) bark lectin recognizes the Neu5Ac(alpha 2-6)Gal/GalNAc sequence. Journal of Biological Chemistry 262, 1596–1601.

Subramanian A, Tamayo P, Mootha VK, Mukherjee S, Ebert BL, Gillette MA, Paulovich A, Pomeroy SL, Golub TR, Lander ES & Mesirov JP (2005) Gene set enrichment analysis: A knowledge-based approach for interpreting genome-wide expression profiles. Proceedings of the National Academy of Sciences 102, 15545–15550.

Tateno H, Toyota M, Saito S, Onuma Y, Ito Y, Hiemori K, Fukumura M, Matsushima A, Nakanishi M, Ohnuma K, Akutsu H, Umezawa A, Horimoto K, Hirabayashi J & Asashima M (2011) Glycome Diagnosis of Human Induced Pluripotent Stem Cells Using Lectin Microarray*. Journal of Biological Chemistry 286, 20345–20353.

Togayachi A, Angata K & Nishihara S (2026) Glycan-related genes and genetic disorders. J Hum Genet, 1–21.

Tu H, Hsiao Y, Yang W, Tsai S, Lin H, Liao C, Lu J, Chou Y, Wang H & Yuh C (2017) Up-regulation of golgi α-mannosidase IA and down-regulation of golgi α-mannosidase IC activates unfolded protein response during hepatocarcinogenesis. Hepatol Commun 1, 230–247.

Uematsu R, Shinohara Y, Nakagawa H, Kurogochi M, Furukawa J, Miura Y, Akiyama M, Shimizu H & Nishimura S-I (2009) Glycosylation Specific for Adhesion Molecules in Epidermis and Its Receptor Revealed by Glycoform-focused Reverse Genomics. Molecular & Cellular Proteomics 8, 232–244.

Varki A (2017) Biological roles of glycans. Glycobiology 27, 3–49.

Wan H, Stone MG, Simpson C, Reynolds LE, Marshall JF, Hart IR, Hodivala-Dilke KM & Eady RAJ (2003) Desmosomal proteins, including desmoglein 3, serve as novel negative markers for epidermal stem cell-containing population of keratinocytes. J Cell Sci 116, 4239–4248.

Wang Y, Khan A, Antonopoulos A, Bouché L, Buckley CD, Filer A, Raza K, Li K-P, Tolusso B, Gremese E, Kurowska-Stolarska M, Alivernini S, Dell A, Haslam SM & Pineda MA (2021) Loss of α2-6 sialylation promotes the transformation of synovial fibroblasts into a pro-inflammatory phenotype in arthritis. Nat Commun 12, 2343.

Wang Y-C, Stein JW, Lynch CL, Tran HT, Lee C-Y, Coleman R, Hatch A, Antontsev VG, Chy HS, O’Brien CM, Murthy SK, Laslett AL, Peterson SE & Loring JF (2015) Glycosyltransferase ST6GAL1 contributes to the regulation of pluripotency in human pluripotent stem cells. Scientific Reports 5, 13317.

Weninger W, Rendl M, Mildner M, Mayer C, Ban J, Geusau A, Bayer G, Tanew A, Majdic O & Tschachler E (2000) Keratinocytes Express the CD146 (Muc18/S-Endo) Antigen in Tissue Culture and During Inflammatory Skin Diseases1. Journal of Investigative Dermatology 115, 219–224.

Yasukawa Z, Sato C & Kitajima K (2005) Inflammation-dependent changes in α2,3-, α2,6-, and α2,8-sialic acid glycotopes on serum glycoproteins in mice. Glycobiology 15, 827–837.

Zhang H, Tsui CK, Castillo JG, Kim EJY, Evangelista A, Liu H, Joe LK, Twells N, Robey EA, Mahal LK, DuPage M & Dillin A (2025) Age-related remodeling of the sialoglycans dampens murine CD8+ T cell function. Science Advances 11, eadw6755.

Zhong J, Liang L, Wu M, Liang Y, Li M, Lee C-L & Shu P (2026) ST6GAL1 Is a Functional Regulator of UVA-Induced Photoaging in Human Dermal Fibroblasts. Cells 15, 1497.

Zhu W, Zhou Y, Guo L & Feng S (2024) Biological function of sialic acid and sialylation in human health and disease. Cell Death Discovery 10, 415.

